# Correlation between antimicrobial structural classes and membrane partitioning: Role of emerging lipid packing defects

**DOI:** 10.1101/2023.07.20.549919

**Authors:** S V Sankaran, Roni Saiba, Samapan Sikdar, Satyavani Vemparala

## Abstract

In this study, a combination of bioinformatics and molecular dynamics simulations is employed to investigate the partitioning behavior of different classes of antimicrobial peptides (AMPs) into model membranes. The main objective is to identify any correlations between the structural characteristics of AMPs and their membrane partitioning mechanisms. The simulation results reveal distinct membrane interactions among the various structural classes of AMPs, particularly in relation to the generation and subsequent interaction with lipid packing defects. Notably, AMPs with a structure-less coil conformation generate a higher number of deep and shallow defects, which are larger in size compared to other classes of AMPs. AMPs with helical component demonstrated the deepest insertion into the membrane. On the other hand, AMPs with a significant percentage of beta sheets tend to adsorb onto the membrane surface, suggesting a potentially distinct partitioning mechanism attributed to their structural rigidity. These findings highlight the diverse membrane interactions and partitioning mechanisms exhibited by different structural classes of AMPs.

## I. INTRODUCTION

Antimicrobial peptides (AMPs) are short chains of amino acids, typically consisting of 12 to 50 amino acids and weighing less than 10 kilodaltons (kDa). These molecules are characterized by their cationic (positively charged) and amphipathic (containing both hydrophobic and hydrophilic regions) nature. Unlike conventional antimicrobial agents, AMPs employ alternative mechanisms to inhibit microbial growth^1^. AMPs are present in various organisms across different classes of life and are often found as secondary metabolites in tissues and mucous membranes. Many of these peptides exhibit a broad range of antibacterial activity, enabling them to effectively eliminate diverse microorganisms^2,3^. Due to their evolutionary conservation in genomes, AMPs are considered essential components of the host defense system against microbial threats^4–8^. AMPs can be categorized based on their source, secondary structure, and amino acid composition^9–11^. They can be derived from various sources such as plants, amphibians, insects, microorganisms, aquatic organisms, and mammals. The presence of a secondary structure, particularly helices, is a crucial characteristic of AMPs. However, there are AMPs that exhibit secondary structures other than helices, including beta strands/sheets, a combination of helices and strands, or even coil structures lacking a distinct secondary structure. Another classification criterion for AMPs is the amino acid composition, which categorizes them as proline-rich, histidine-rich, tryptophan-rich, glycine-rich, or arginine-rich peptides^9^. The primary mode of action involves disrupting the integrity of microbial membranes, although they can also exert intracellular functions^2^.

AMPs have the ability to interact with membranes even at low concentrations in their monomeric state, and these interactions can significantly impact membrane properties. This ranges from thinning of the membrane due to partial partitioning of the peptides to complete translocation into the cellular environment. However, the precise mechanisms underlying the antimicrobial action of AMPs are not fully understood and are thought to involve several processes, including sensing, partitioning, disruption, and eventual cell lysis. The key components of many AMPs are charged and hydrophobic residues, which have been implicated in the initial sensing and differentiation of bacterial membranes from host membranes, as well as in subsequent partitioning processes. However, it has become increasingly clear that this simplistic view may not capture the complete picture. Recent studies have revealed that AMPs can sense not only differences in lipid components with opposite charges but also intrinsic membrane properties such as lipid packing defects and curvature as part of their mechanism to differentiate between bacterial and host cell membranes. In addition to recognizing specific lipid compositions, AMPs can sense and respond to variations in the structure and organization of membranes. Lipid membranes possess inherent characteristics related to the presence of packing defects, which are influenced by the types and compositions of lipids. The dynamics occurring at the interface between lipids and water can cause random protrusions of hydrophobic lipid tails into the surrounding solvent, resulting in the formation of these packing defects^12–15^. For example, bacterial membranes are known to exhibit higher levels of lipid packing defects compared to host membranes. These packing defects, which result from the stochastic dynamics of lipid molecules at the lipid-water interface, provide cues that enable AMPs to distinguish bacterial membranes. Moreover, differences in membrane curvature between bacterial and host cells have also been shown to contribute to the antimicrobial mechanism of AMPs. These findings highlight the complexity of membrane recognition by AMPs and suggest that a combination of factors, including lipid composition, packing defects, and curvature, may contribute to their antimicrobial activity. Further research is necessary to elucidate the interplay between these different mechanisms and to fully understand the umbrella of antimicrobial action employed by AMPs. Over the past few years, several research groups have contributed to the understanding and characterization of these lipid packing defects^12,13,16–20^. Different methodologies have been employed, including a solvent accessible surface area (SASA)-based method developed by Voth and colleagues^12^, as well as a Cartesian grid-based approach used by other researchers to scan the lipid-water interface and identify solvent-exposed lipid atoms associated with packing defects^13,14^. The latter method has been incorporated into the Packmem software^21^, which utilizes an algorithm that divides the x-y plane of the lipid bilayer into grids and scans along the z-direction from the solvent interface down to a level below the average position of the C2 atoms of the glycerol moieties. This scanning process allows the identification of regions with low lipid density, and the packing defect sites are characterized quantitatively based on their occupied area (A) and qualitatively labeled as ”Deep” or ”Shallow” relative to the average level of the C2 atoms. While this method primarily focuses on projecting the defect area onto the membrane’s x-y plane, Tripathy et al.^17^ introduced a quantitative approach to assess both the depth and area of the defects. They achieved this by scanning the local free volume surrounding each lipid atom to identify defect pockets within the membrane.

The primary objective of this paper is to uncover universal principles governing the initial interaction between structural classes of antimicrobial peptides (AMPs) and bacterial membranes, with a specific focus on lipid packing defects. To achieve this goal, the following approach is undertaken:

1. Database analysis: An existing database of AMPs is utilized to identify sequences for which structural information is available. This subset of AMPs is then classified into different structural classes based on specific variables or criteria.
2. Selection of representative AMPs: From the identified structural classes, representative AMPs are chosen to ensure a diverse representation of structural characteristics.
3. Molecular dynamics simulations: Molecular dynamics simulations are conducted for the selected representative AMPs in solution and in the presence of model bacterial membranes. These simulations provide insights into the dynamic behavior and interactions of AMPs with membranes at an atomic level.

## II. MODELS AND METHODS

A flowchart describing the various steps in the pipeline of data collection, classification of structural classes. and identification of representative AMPs for MD simulations is illustrated in Fig. 1.

**FIG. 1.**
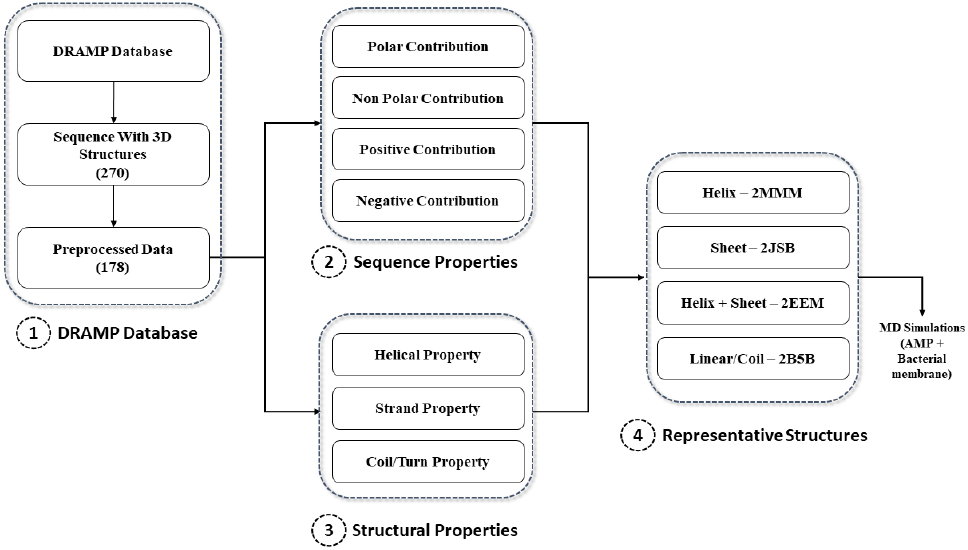
Analyses and simulation pipeline used in this study

### A. Analysis of DRAMP database

DRAMP^22,23^ (Data Repository of AntiMicrobial Peptides) is a manually curated open-source database consisting annotations for diverse set of AMPs including their sequences, structures, activities, literature references, clinical and physiochemical information. Currently, the database harbours ∼25000 entries, of which 6105 are general AMPs (both synthetic and natural), 16110 patented AMPs and 77 AMPs that are under preclinical trials. Based on the activity of the peptide from the literatures, the database has grouped the compounds into 11 different classes such as antibacterial, antifungal, etc. This database can be accessed using the specified URL http://dramp.cpu-bioinfor.org/

### B. MD simulation in model bacterial Membrane

To model the bacterial membrane, we use a pre-equilibrated membrane patch consisting of 70% POPE (palmitoyl-oleoyl-phosphatidylethanolamine) and 30% POPG (palmitoyl-oleoyl-phosphatidylglycerol) lipids. This membrane composition closely resembles the inner membrane of bacteria^24^. The membrane patch, containing 128 lipids per leaflet, is prepared using the Membrane Builder module of CHARMM-GUI^25^, which has been utilized in our previous studies on antimicrobial polymers^26–28^. For each antimicrobial peptide (AMP), we select a random conformation from the solution state NMR derived structural ensemble. The protonation states of the peptide residues are determined at neutral pH using PROPKA3^29,30^, and hydrogen atoms are added accordingly. To set up the four AMP-membrane systems, the peptides are placed near the upper leaflet of the pre-equilibrated POPE-PG bilayer, ensuring there are no steric clashes. We add sufficient water molecules and ions to maintain a salt concentration of 0.15 M. The all-atom molecular dynamics (MD) simulations of the systems are performed using the TIP3P water model^31^, along with CHARMM36m^32^ and CHARMM36^33^ force field parameters for peptide and lipid molecules, respectively. The simulations are conducted using NAMD2.10^34^. Prior to the production run, all systems undergo an energy minimization of 10,000 steps. Then, an equilibration protocol is applied, where positional restraints on peptide heavy atoms are gradually decreased over a period of 3 nanoseconds to ensure relaxed starting configurations of the AMP-membrane systems. The simulations are performed under periodic boundary conditions in the isothermal-isobaric ensemble at 1 atm pressure and 310 K, with a time step of 2 femtoseconds. van der Waals interactions beyond 12Åare smoothly truncated using a force-based switching function, while electrostatic interactions are calculated using the Particle mesh Ewald fast Fourier transform. The production runs are carried out for over 300 ns, and subsequent analysis of the equilibrated trajectories is performed using VMD^35^, MEMBPLUGIN^36^, and Packmem^21^. The MD system details can be found in Table. S1.

## III. RESULTS

### A. Identification of Structural Classes

#### 1. Dataset filtration

A dataset containing 4,039 antibacterial sequences is obtained from the DRAMP database^22,23^. Each sequence in the dataset is associated with various attributes such as name, SWISSPORT entry, family, gene, source, activity, protein existence, structure, structure description, PDB ID, Binding Target, PubMed ID, references, etc. From this dataset, we identify 230 sequences for which 3D structural data is available in the Protein Data Bank^37^. However, some sequences are derived from larger proteins, and the corresponding PDB structures contain the entire protein rather than just the antimicrobial peptide (AMP). To rectify this issue, we manually examine the dataset and select a set of 178 sequences that have 3D structures specifically for the peptides, excluding the larger protein structures. Properties such as hydrophobicity, net charge, and amphipathicity are crucial for AMPs^38,39^, and these properties are calculated based on the amino acid side chains of the selected sequences. For each sequence, the counts of each property are obtained by classifying the amino acid side chains. To assess the contribution of each property in a given sequence, the counts are normalized based on the sequence length. The contribution estimates the significance of each property in the sequence. In addition to sequence-level properties, the secondary structure (SS) content of AMPs plays a vital role in their mechanism of action^40^. The presence of helical structure, for example, is important for maintaining the amphipathic nature of the peptide, allowing it to interact with both the hydrophilic head and the hydrophobic tail of the membrane. The secondary structure elements, such as helix, sheet, coil, and turn, are determined using the STRIDE algorithm^41^ implemented in Visual Molecular Dynamics (VMD)^35^. Similar to the physicochemical properties, the secondary structural contents are normalized with respect to the sequence length to enable comparative analysis.

To gain insights into the relationship between structure and sequence properties, an exploratory analysis is conducted on the physicochemical and structural properties of antimicrobial peptides (AMPs). Firstly, a histogram is generated to visualize the distribution of sequence lengths, with a bin width of 5 (Supplementary Information, SI, Fig. S1). The analysis reveals that the sequence lengths of AMPs range from a minimum of 9 to a maximum of 93. While the range is broad, the majority of peptides have lengths between 10 and 45, indicating that AMPs commonly fall within this range. Box plots are then used to depict the percentage contribution of sequence properties (Fig. S2). The results indicate that, on average, AMP sequences consist of approximately 40 to 60% non-polar residues, 10 to 40% polar residues, 15 to 30% positively charged residues, and up to 10% negatively charged residues. These box plots provide valuable insights into the amino acid composition of AMPs, highlighting the need for a balanced distribution of different types of residues. Achieving the right balance is crucial as it directly influences the physicochemical properties of the peptides, which in turn play a crucial role in their antimicrobial activity.

Based on the analysis of physicochemical and structural properties, we have identified four representative AMPs that belong to different secondary structural classes^38^. These classes include *α*-helix, *β*-sheet, *α*-helix+*β*-sheet, and disordered/coil structures. The AMPs chosen as representatives are Aedesin (*PDB id: 2MMM*) with an *α*-helical structure^42^, Arenicin-1 (*PDB id: 2JSB*) with a *β*-sheet structure^43^, Mytilin (*PDB id: 2EEM*) with an *α*-helix+*β*-sheet structure^44^, and Tewp (*PDB id: 2B5B*) with a disordered/coil structure^45^. These peptides will be referred to as *α*-peptide, *β*-peptide, *α* +*β*-peptide, and coil-peptide, respectively. To investigate the effects of secondary structure on the partitioning mechanisms of these AMPs, all-atom molecular dynamics (MD) simulations are performed in the presence of a model bacterial membrane. The distribution of secondary structural properties, such as helix, strand, coil, and turn, is studied (Figure 2A). The analysis reveals that AMPs are designed with a preference for a helical secondary structure over other secondary structures. It is worth noting that out of the 178 structures, 8 are completely coil in nature. The chosen four representative peptides, based on their secondary structural contributions and sequence length can be seen in Fig. 3. The chosen representatives have the highest contributions to their respective secondary structures. However, for the Helix+strand representative, both secondary structures contribute equally. Except for the strand representative, all other representatives have sequence lengths ranging from 34 to 36. The significant difference in sequence length for the strand representative (21) is due to the observed negative correlation between sequence length and strand percentage.

**FIG. 2.**
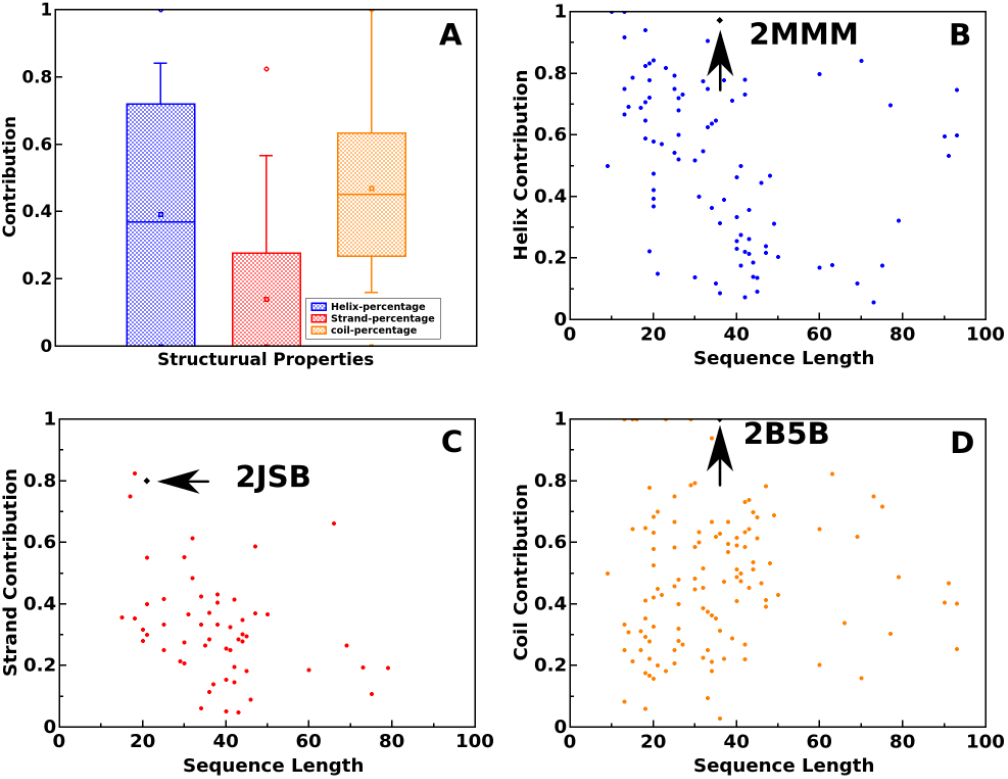
A) The box plot showing the contribution from all the 3 structural properties. B) Sequence length vs helix contribution. C) Sequence length vs strand contribution. D) Sequence length vs coil contribution

**FIG. 3.**
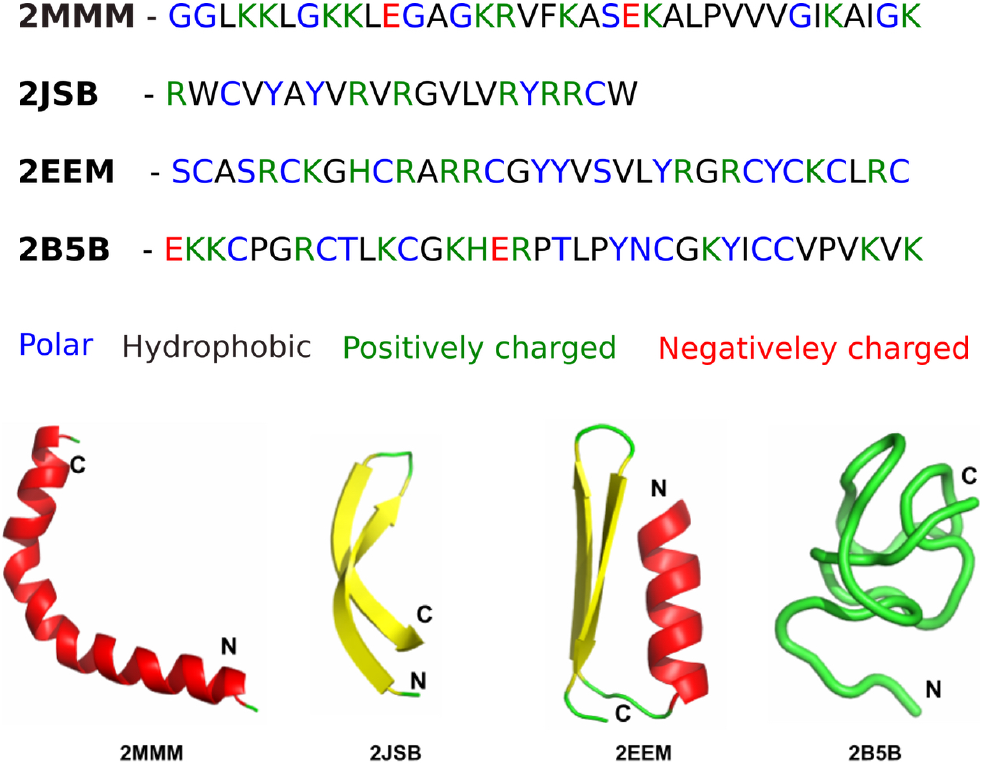
Amino acid sequence from 4 different classes of the peptides colour coded based on the properties. Blue - Polar; red - Negatively charged; Green - positively charged; black - hydrophobic. Representative classes of peptides chosen based on DRAMP database analysis. The structures are coloured based on their secondary structure: helix (red), *β* sheet (yellow), coil/turn (green)

### B. Partitioning of AMPs into model membranes

In this section, we aim to quantify the interactions between the representative AMPs and the model bacterial membrane, focusing on both structural changes to the peptides and the membrane itself. The initial setup of AMPs near the model membrane can be seen in Fig. S3 and Fig. 4 displays snapshots of the peptides and membrane at the end of the simulation runs. All four peptides exhibit varying degrees of partitioning into the model bacterial membranes, with the *α*-peptide displaying the deepest insertion within the simulation time scale considered in this study. To assess the structural changes of the peptides, we measure the root mean squared deviation (RMSD) and radius of gyration (*R*_*g*_), which are depicted in Fig. S4 (see Supplementary Information). The *α*-peptide exhibits the greatest deviation from the initial crystal structure during its interaction with the membrane. These structural changes are further reflected in the variations of *R*_*g*_ and the evolution of secondary structure of the peptides, as illustrated in Figs. S4 and S5 (see Supplementary Information). Specifically, for the *α*-peptide, as the simulation progresses and the peptide-membrane interaction intensifies, a noticeable loss of *α*-helical content is observed (Fig. S5(a)). In the case of the *α*+*β*-peptide, clear indications of decreased *α*-helical and *β*-sheet content are observed as the interaction with the membrane increases. However, no significant changes in the secondary structure content are observed for the *β*- and coil-peptides. To gain further insights into the dynamics of peptide insertion into the membrane environment, we tracked the time evolution of the *z*-coordinate trajectories of key hydrophobic residues in each peptide, as depicted in Fig. 5 and hydrophilic residues in Fig. S6. For the *α*-peptide, the hydrophobic residues exhibit coordinated insertion, where the partitioning of one hydrophobic residue facilitates the insertion of other hydrophobic residues. A similar trend is observed for the *β*-peptide insertion. In contrast, the insertions in the *α* + *β*-peptide and coil-peptide are not coordinated. This discrepancy may be attributed to the initial coil content of these peptides. The presence of significant coil content allows for conformational malleability, wherein the absence of structural requirements imposed by rigid secondary structure elements (present initially) enables the independent partitioning of residues. The insertion depth of various parts of the AMPs considered are mapped on to the structural elements of the AMPs and is shown in Fig. 6. The relevant hydrophobic residues are also shown. The insertion of hydrophobic residues into the membrane can also have profound effects on lipid packing defects that are sensed and exploited by membrane-active peptides. These effects will be further explored in the subsequent section.

**FIG. 4.**
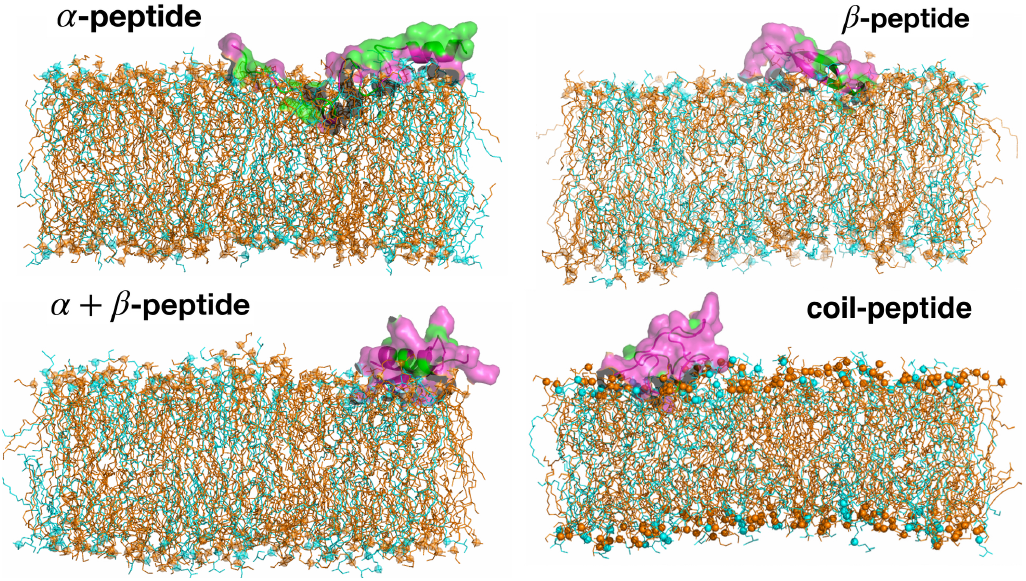
Snapshots of partitioning of four AMPs into a model bacterial along the membrane normal, at the end of the simulation runs, are shown. Water and ions are not shown for clarity. The hydrophobic and hydrophilic residues of AMPs are coloured in green and pink respectively. The POPE and POPG lipid molecules in the membrane are coloured as orange and cyan respectively.

**FIG. 5.**
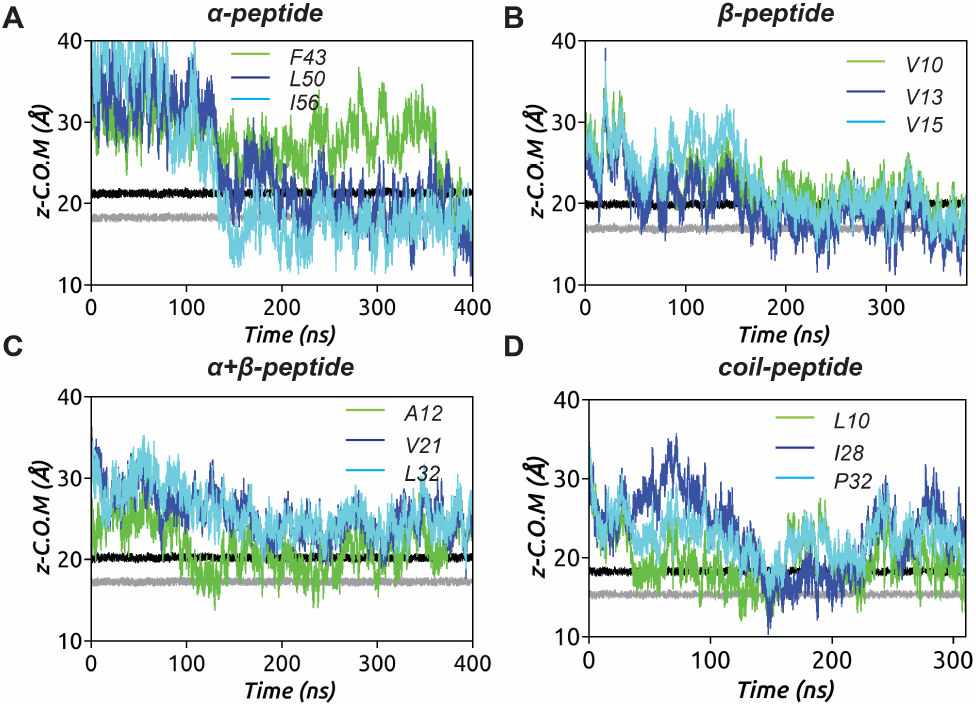
Partitioning of key hydrophobic residues into the model bacterial membrane during the simulation for four AMPs considered. The locations of lipid headgroup P atoms (black) and C2 atoms (grey) of glycerol moities of the upper leaflet, along the membrane normal, of the model bacterial membrane are also shown.

**FIG. 6.**
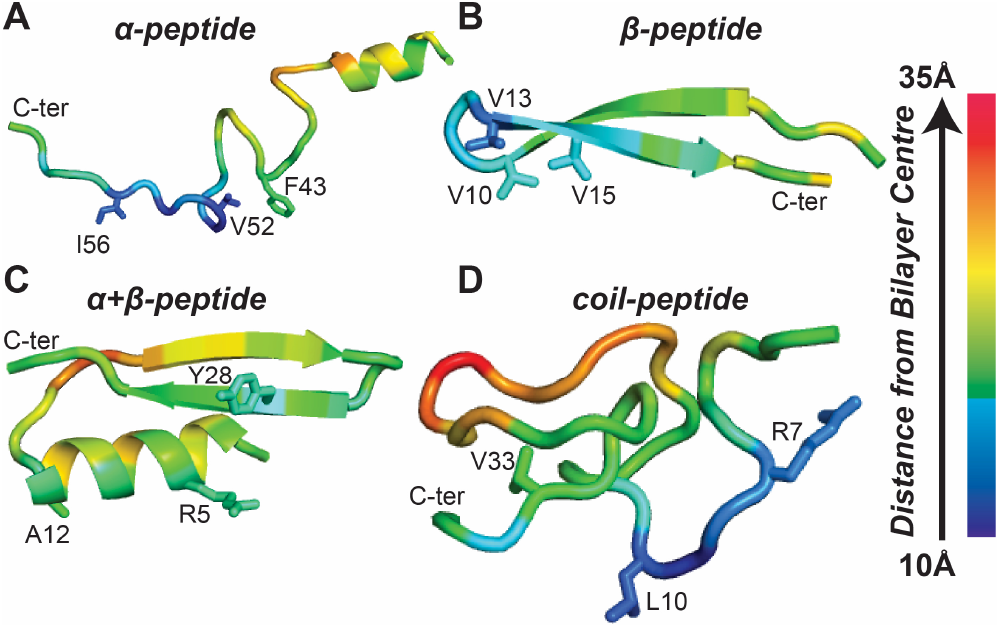
The average residue insertion depths of the four AMPs are calculated as distance from bilayer centre averaged over last 50 ns. The colour scale indicates, the smaller the value, the deeper is the insertion.

To analyze the emergence of amphiphilicity in the partitioned peptides, *z*-density profiles of the four peptides are calculated over the last 100 ns of the simulation. The density profiles, shown in Fig. 7, provide insights into the distribution of hydrophobic and hydrophilic residues in the peptides within the membrane. By comparing the peak positions of the hydrophobic and hydrophilic density profiles, we can estimate the amphiphilicity content of the peptides. The data indicates that the *α*-peptide exhibits the highest level of amphiphilic character, despite undergoing significant structural changes as observed in Fig. S5(a). The location of the hydrophobic residue density peak further illustrates the extent of hydrophobic residue partitioning in the *α*-peptide. In the case of the *α*-peptide, which demonstrates the deepest insertion into the membrane environment, we can identify four initial stages of the process. First, in the recognition stage, the AMP detects the presence of the bacterial membrane through electrostatic interactions and possibly by sensing lipid packing defects (further discussed in the next section). Next, the AMP adheres to the membrane surface and adapts to the environment, which may lead to structural changes. As a result of this adaptation, the *α*-peptide partitions into the interior of the membrane. This behavior is effectively captured by the hydrophobic moment vector of the *α*-peptide, as illustrated in Fig. 8.

**FIG. 7.**
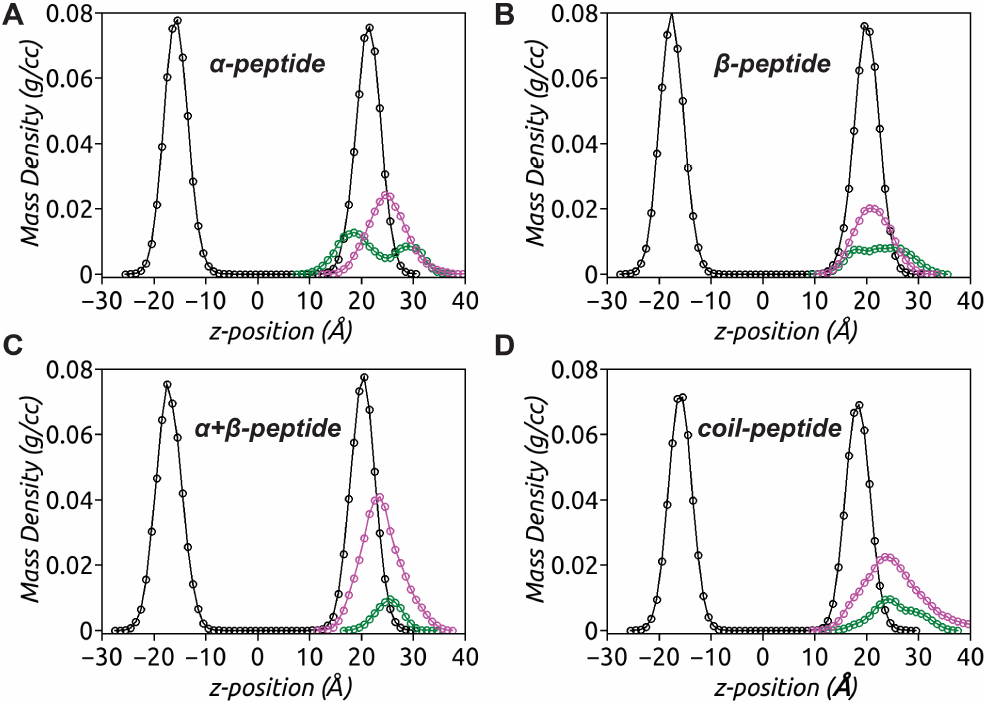
The z-density profiles of the four AMPs interacting with model bacterial membrane indicate the location and extent of hydrophobic (green) and hydrophilic (magenta) density components of the peptide with respect to lipid head-group phosphate atom (P) density (black), calculated over the last 100 ns of respective MD trajectories.

**FIG. 8.**
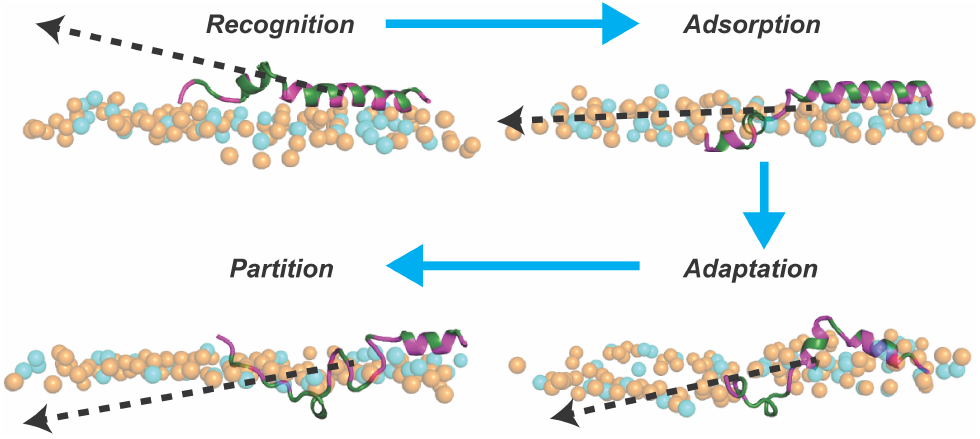
The different stages of *α*-peptide interacting with a model bacterial membrane composed of POPE (orange) and POPG(cyan) lipids is illustrated. The peptide-membrane recognition is driven by electrostatic interactions between hydrophilic (magenta) peptide residues and lipid headgroups, followed by adsorption on the membrane surface. The unfolding of *α*-peptide favors conformational adaptation such that hydrophobic (green) residues face membrane interior, while the hydrophilic ones reside close to lipid headgroups, subsequently gaining facial amphiphilicity, resulting in peptide partitioning. The change in orientation of the 3d-Hydrophobic Moment (3d-HM) vector (black dotted arrow) from initially being away from the membrane surface to finally facing the membrane interior demonstrates how *α*-peptide unfolding facilitates access of hydrophobic residues to the membrane core.

### C. Distribution of defect size

In this section, we investigate how different classes of AMPs modulate and influence lipid packing defects in model bacterial membranes. The analysis of lipid packing defects is conducted using the Packmem software^21^ over the last 200 ns of each system. To characterize the abundance of defect sites per leaflet in a given frame, denoted as *Nsites*, we examine the distributions *P* (*Nsites*) separately for deep and shallow defects. The results are presented in Fig.10A and Fig.10B, respectively. The average number of deep defect sites per leaflet (*< Nsites >∼* 32) is found to be similar for the *α*-, *β*-, and *α* + *β*-peptides, as indicated by the overlapping distributions in Fig. 10A. However, the coil-peptide induces a higher abundance of deep defect sites (*< Nsites > ∼* 42) compared to the structured AMPs. A similar trend is observed for the distributions of shallow defect sites in Fig. 10B. The coil-peptide leads to a greater number of shallow defects compared to the structured AMPs. Overall, these results suggest that the coil-peptide, with its disordered nature, engages in larger surface contacts with the bilayer, leading to an increased number of lipid packing defects compared to the structured AMPs.

The size distribution, denoted as *P* (*A*), of defect sites is an important aspect to characterize the effects of structurally diverse AMPs. The defect size distributions, represented in semi-log scale as *log*_10_*P* (*A*), are illustrated in Fig.10C and Fig.10D. It is interesting to observe that even though the number of deep defect sites is similar across the three structured peptides, the size distributions of deep defects exhibit significant differences (Fig. 10C). The coil-peptide shows a high population of moderate-sized deep defects (∼ 40 − 70Å^2^), followed by the *α*-peptide, *β*-peptide, and *α* + *β*-peptide. However, larger deep defect sites (*>* 80Å^2^) are more pronounced in the presence of the *α*-peptide, particularly due to the insertion of several hydrophobic residues into a single colocalized large deep defect site. The observed pattern in deep defect size distributions can be attributed to the predominance of hydrophobic residues in the *α*-peptide, while the *α* + *β*-peptide has the least number of such residues. In contrast, the size population of shallow defect sites demonstrates that the coil-peptide not only enhances the number but also the size of shallow defects compared to the structured peptides, which exhibit similar distributions (Fig. 10D). This in particular is also evident from membrane response in terms of the corresponding 2D-thickness maps generated for the final snapshots of AMP-membrane systems (Fig. 9). It is clearly evident that owing to abundance of defect sites in presence of the coil-peptide, the model bacterial membrane exhibits global thinning. However, membrane thinning is only locally induced at the *α*-peptide insertion site. In contrast, membrane thinning is compromised in presence of the other two structural classes of AMPs consistent with their observed degree of partitioning and induced packing defect sites.

**FIG. 9.**
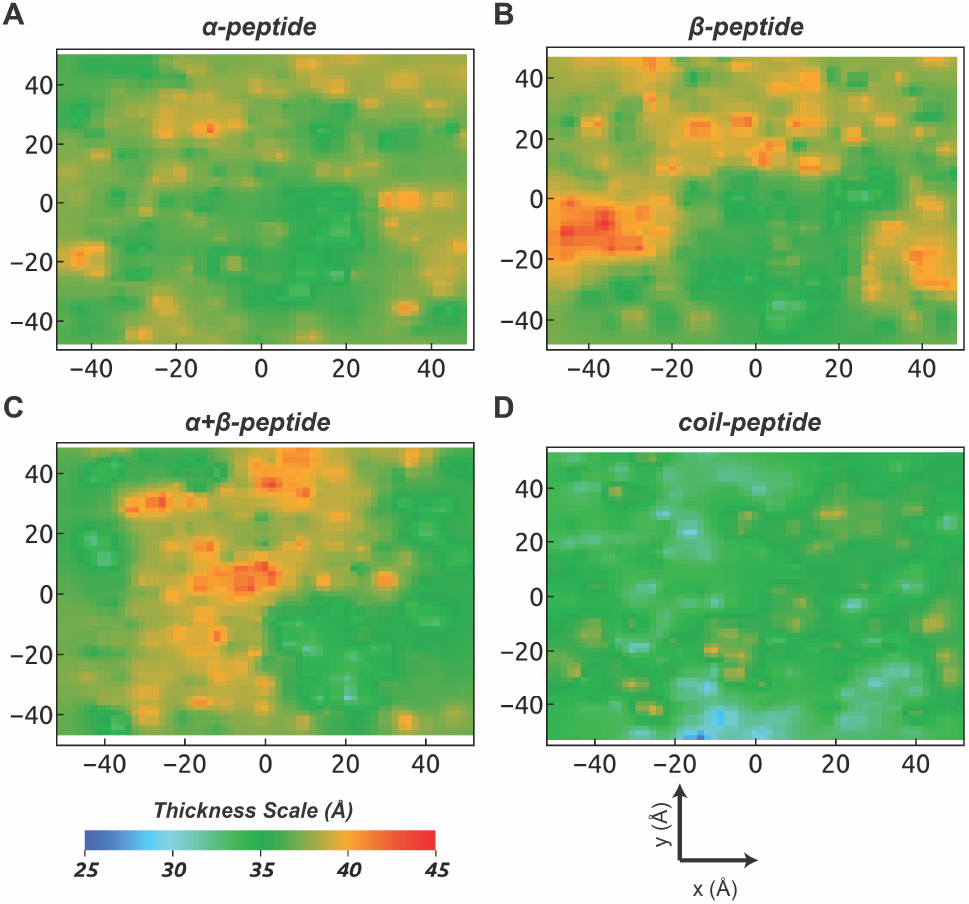
The 2D-thickness maps of model bacterial membrane in presence of the four AMPs. The thickness maps based on inter-leaflet P-P distance are generated using a 2 Å resolution along the x-y plane for the final MD snapshots of respective trajectories.

**FIG. 10.**
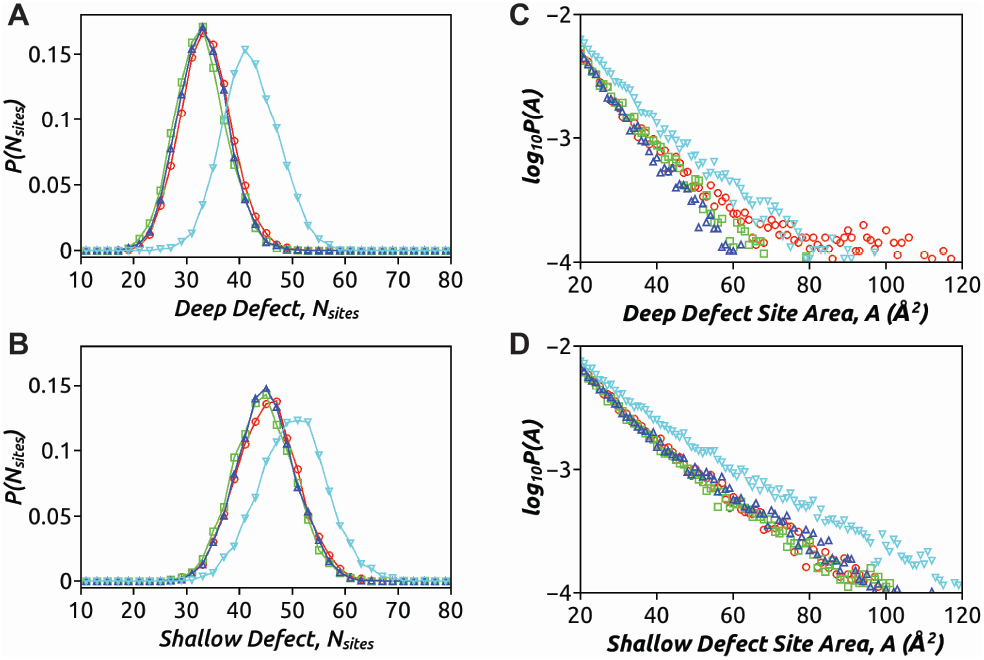
The distribution of number of defect sites per leaflet in a given frame, *P* (*Nsites*), for (A) deep and (B) shallow defects. The size distribution of defect sites, *log*10*P* (*A*), for (C) deep and (D) shallow defects. The analysis is performed over the last 200 ns trajectories of each AMP-membrane system:*α*-peptide (ellipse, red), *β*-peptide (square, green), *α*+*β*-peptide (up-triangle, blue) and coil-peptide (down-triangle, cyan)

In this section, we illustrate a cooperative interplay involving the appearance of a defect site on the membrane surface co-localized with the *α*-peptide (Fig.11). As mentioned earlier, the C-terminal unfolding of the *α*-peptide precedes the hydrophobic residue insertions. We consider the center of mass of these hydrophobic residues from the unfolded helix and simultaneously track the insertion dynamics using the *z*-distance of this center of mass from the average level of C2 atoms in POPE-PG lipids along the membrane normal (z-direction). We also monitor the appearance of any underlying packing defects on the membrane surface (Fig.11A). It is observed that at around 150 ns, a large co-localized deep defect arises, favoring the transient insertion of the unfolded part of the helix, as indicated by negative values of the z-distance. This co-localized deep defect site continues to grow over time, facilitating discrete events of hydrophobic residue insertions and resulting in the partitioning of the *α*-peptide. This process is characterized by the sensing of lipid packing defects. The final snapshot illustrating the insertion of the *α*-peptide into this co-localized deep defect site is shown in Fig. 11B. The location of defect sites for the other AMPs is shown in Supplementary Information, Fig. S7.

**FIG. 11.**
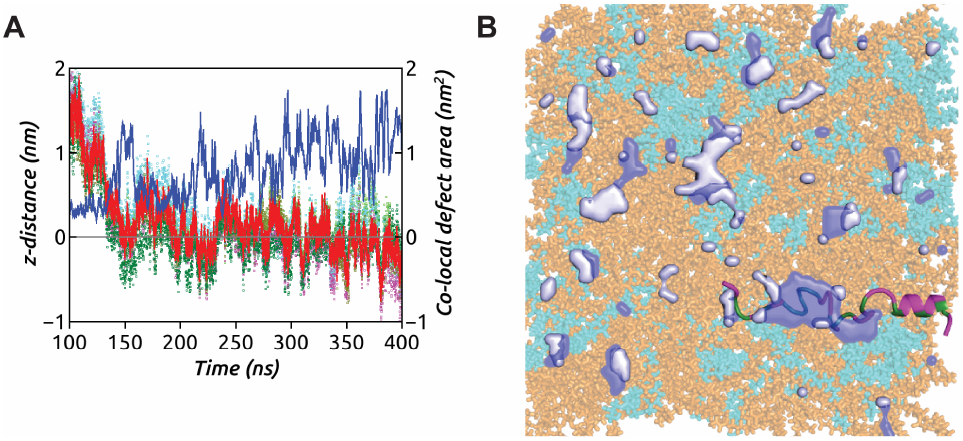
(A) The insertion dynamics of centre of mass of hydrophobic residues (red line) from C-terminal of the *α*-peptide into a co-localized deep defect site, the area fluctuations being shown in blue. Centre of masses of individual hydrophobic residues are indicated in dots of different colours. (B) The final snapshot of *α*-peptide-POPE (orange)/PG (cyan) system illustrates the location and extent of the co-localized deep defect (dark blue) surrounded by shallow defect sites (light blue). The hydrophobic residues of the peptide are shown in green, while the hydrophilic ones in magenta.

## IV. DISCUSSION

The role of lipid packing defects in the recruitment of amphiphilic molecules has been extensively studied in various systems. Hydrophobic nanoparticles, disaccharides, and proteins have been shown to partition into membranes by sensing lipid packing defects^14–16,46–51^. The concept of the amphipathic lipid packing sensor (ALPS) motif, commonly found in peripheral membrane proteins, has shed light on the role of hydrophobic residues in sensing and inserting into lipid packing defects^14–16,52^. Vanni et al. demonstrated that the hydrophobic residues of the ALPS motif act as lipid packing sensors and subsequently insert into pre-existing packing defects^14^. However, a later study suggested that the ALPS motif in close proximity to the membrane can promote the formation of new defects, indicating that the pre-existence of a packing defect is not necessary for peptide insertion^16^.

Recent research has also highlighted the importance of lipid packing defects in regulating the partitioning mechanism of viral peptides and antimicrobial polymers in membranes. For example, in the case of the Hepatitis A virus 2B (HAV-2B) peptide, the presence of lipid packing defects facilitated peptide partitioning, while a cholesterol-rich bilayer with fewer defects inhibited partitioning^18,53^. Another study demonstrated how antimicrobial polymers with different chemical compositions explored and occupied lipid packing defects at various depths, exhibiting different partitioning mechanisms into model bacterial membranes^54^. Investigations by Voth and co-workers have revealed that packing defects increase with an increase in membrane curvature^12^. The synergistic effect of membrane curvature and lipid tail unsaturation on interfacial packing defects has also been confirmed, where increasing lipid unsaturation or introducing conical lipids into flat bilayers led to a defect size distribution resembling that of a positively curved bilayer^15^. Furthermore, studies have shown that membrane thinning enhances lipid packing defects. By applying external force in molecular dynamics simulations, Hilten et al. developed a protocol to vary membrane thickness and found a higher probability of finding large defects in thin membranes compared to membranes of normal thickness^52^.

In this study, a combination of bioinformatics and molecular dynamics simulations is utilized to explore how diverse classes of antimicrobial peptides (AMPs) interact with model membranes and partition within them. The primary objective is to uncover potential correlations between the structural characteristics of AMPs and their mechanisms of membrane partitioning.The simulation results reveal distinct membrane interactions among the various structural classes of AMPs, particularly in relation to the generation and interaction with lipid packing defects. Notably, AMPs with a structure-less coil conformation generate a higher number of deep and shallow defects, which are larger in size compared to other classes of AMPs. Conversely, AMPs with a helical component demonstrate the deepest insertion into the membrane. On the other hand, AMPs with a significant percentage of beta sheets tend to adsorb onto the membrane surface, suggesting a distinct partitioning mechanism due to their structural rigidity. The different classes of AMPs modulate lipid packing defects to varying extents, influenced by factors such as hydrophobic residue insertions, surface area contacts, or a combination of both. This indicates the existence of diverse mechanisms for perturbing bacterial membranes. In the field of designing biomimetic polymers with antimicrobial properties^20,26,27,55–57^, these findings hold significance, as they provide insights into how naturally occurring antimicrobial peptides interact and exploit bacterial membrane lipid packing defects for efficient partitioning. Such knowledge can aid in the better design of antimicrobial polymers. Future studies will also investigate how aggregates of AMPs from different structural classes interact and influence membranes through lipid packing defects, further advancing our understanding of the antimicrobial mechanisms employed by these peptides.

## ACKNOWLEDGEMENT

All the simulations in this work have been carried out on clusters Annapurna and Nandadevi at The Institute of Mathematical Sciences, Chennai, India.

