## Supplemental Information for "Correlation between antimicrobial structural classes and membrane partitioning: Role of emerging lipid packing defects"

(Dated: July 20, 2023)

*
†
‡
§

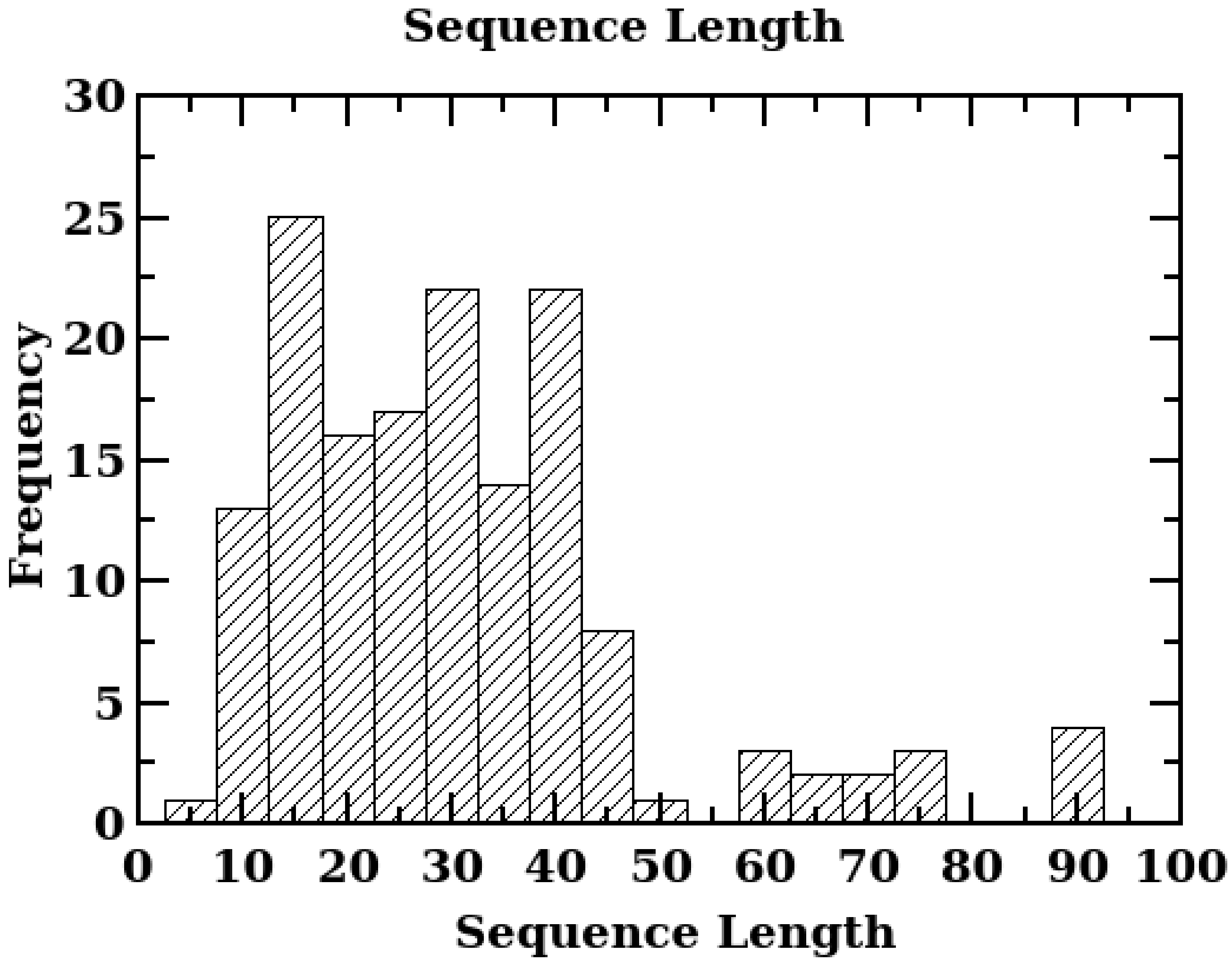


FIG. S.1. Distribution of sequence lengths of AMPs in DRAMP data base.

TABLE S.1. MD system details

| Peptide | Simulation time (ns) | Initial box size (Å$^3$) | Total atoms | Lipid atoms | Water atoms |
|---|---|---|---|---|---|
| 2B5B | 345 | 110 x 110 x 110 | 91379 | 22500 | 58470 |
| 2EEM | 400 | 110 x 110 x 80 | 67044 | 22500 | 43873 |
| 2JSB | 368 | 110 x 110 x 85 | 63313 | 22500 | 30639 |
| 2MMM | 400 | 110 x 110 x 105 | 78979 | 22500 | 46104 |

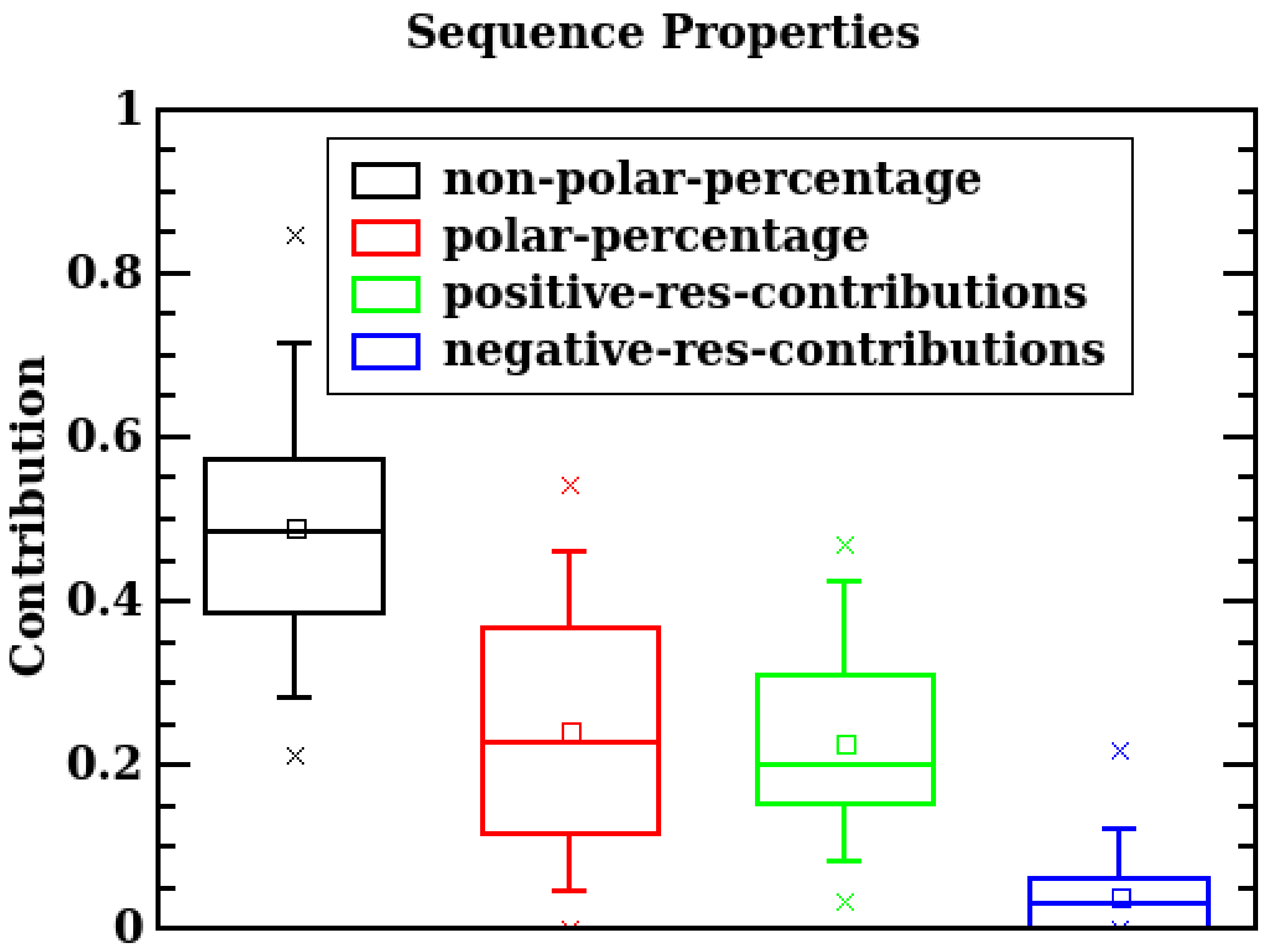


FIG. S.2. Box plot of distribution of different classes of residues of AMPs in DRAMP data base

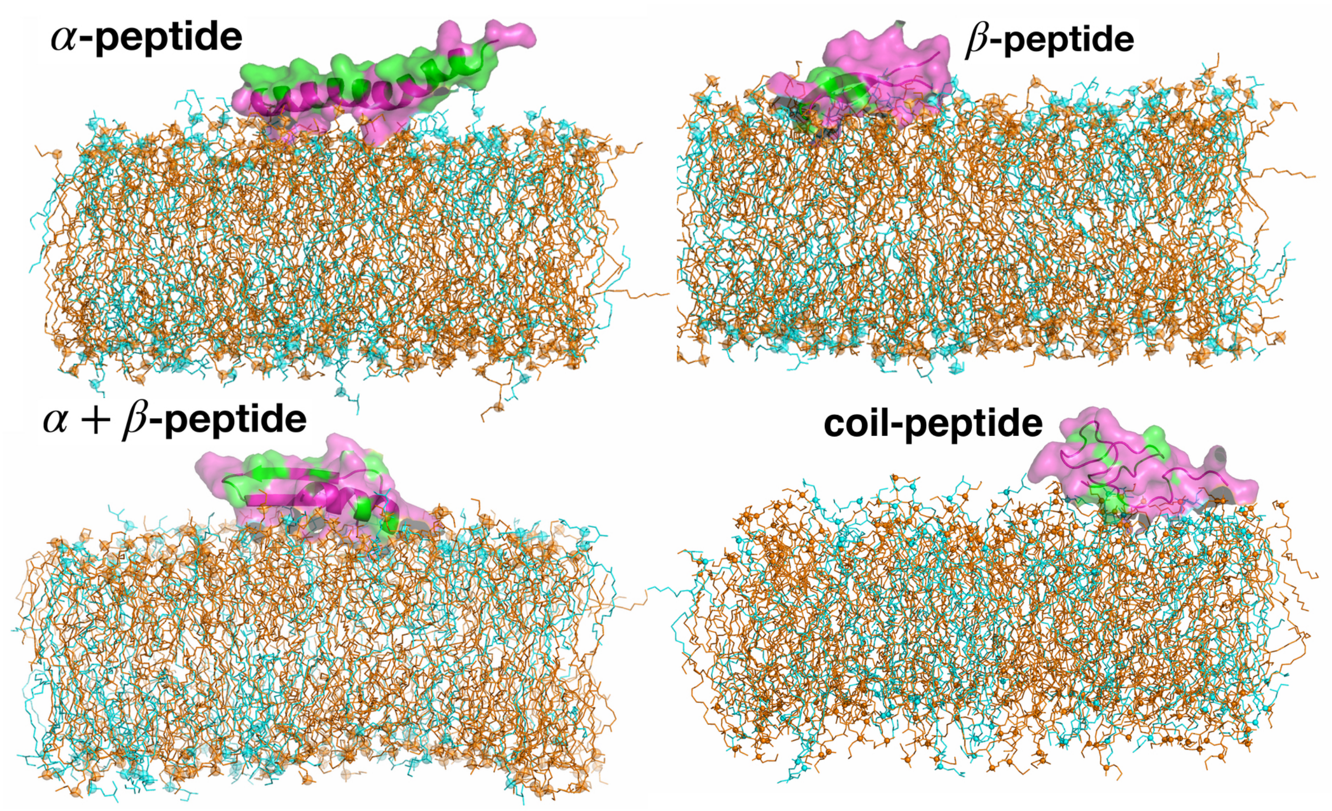


FIG. S.3. Initial setup of four AMPs on a model bacterial along the membrane normal are shown. Water and ions are not shown for clarity. The hydrophobic and hydrophilic residues of AMPs are coloured in green and pink respectively. The POPE and POPG lipid molecules in the membrane are coloured as orange and cyan respectively.

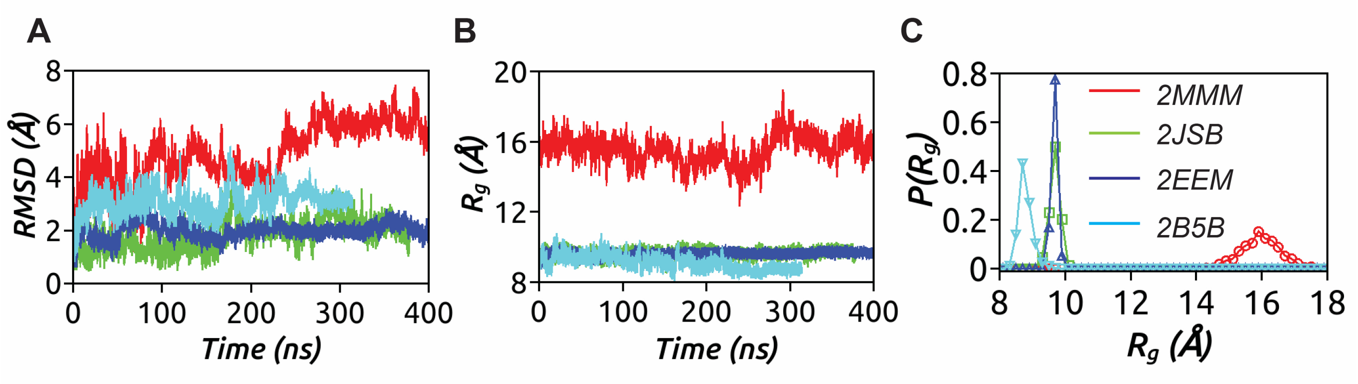


FIG. S.4. Structural properties of simulated four AMPs in the bilayer environment. (a) root mean squared deviation from the initial solution crystal structures (b) radius of gyration indicating the size of the AMPs (c) conformational sampling of the AMPs, in terms of $R_g$, to understand changes in the structures as the membrane simulation progresses.

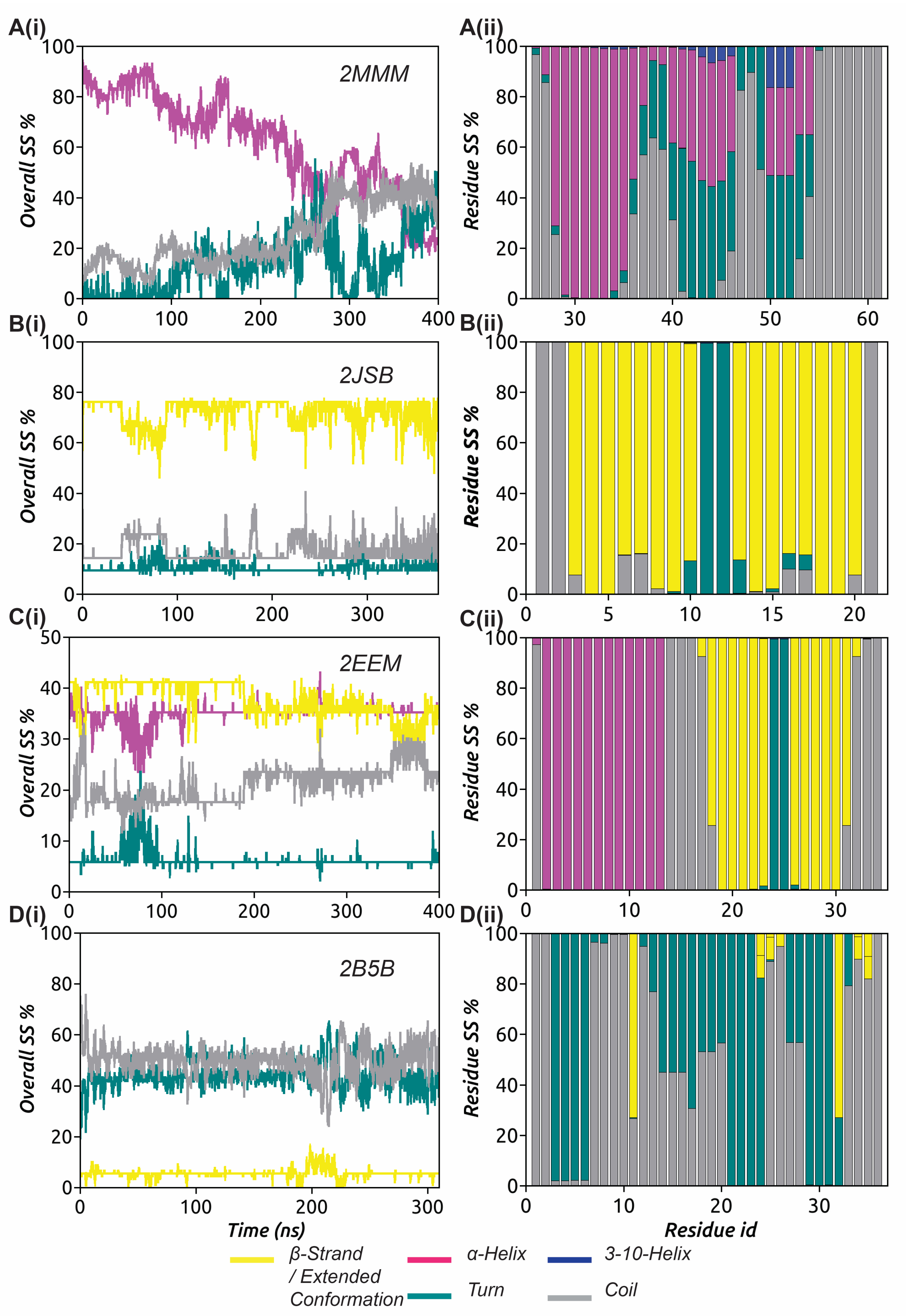


FIG. S.5. Evolution of secondary structural elements in the four AMPs during the interaction with the model bacterial membrane are shown.

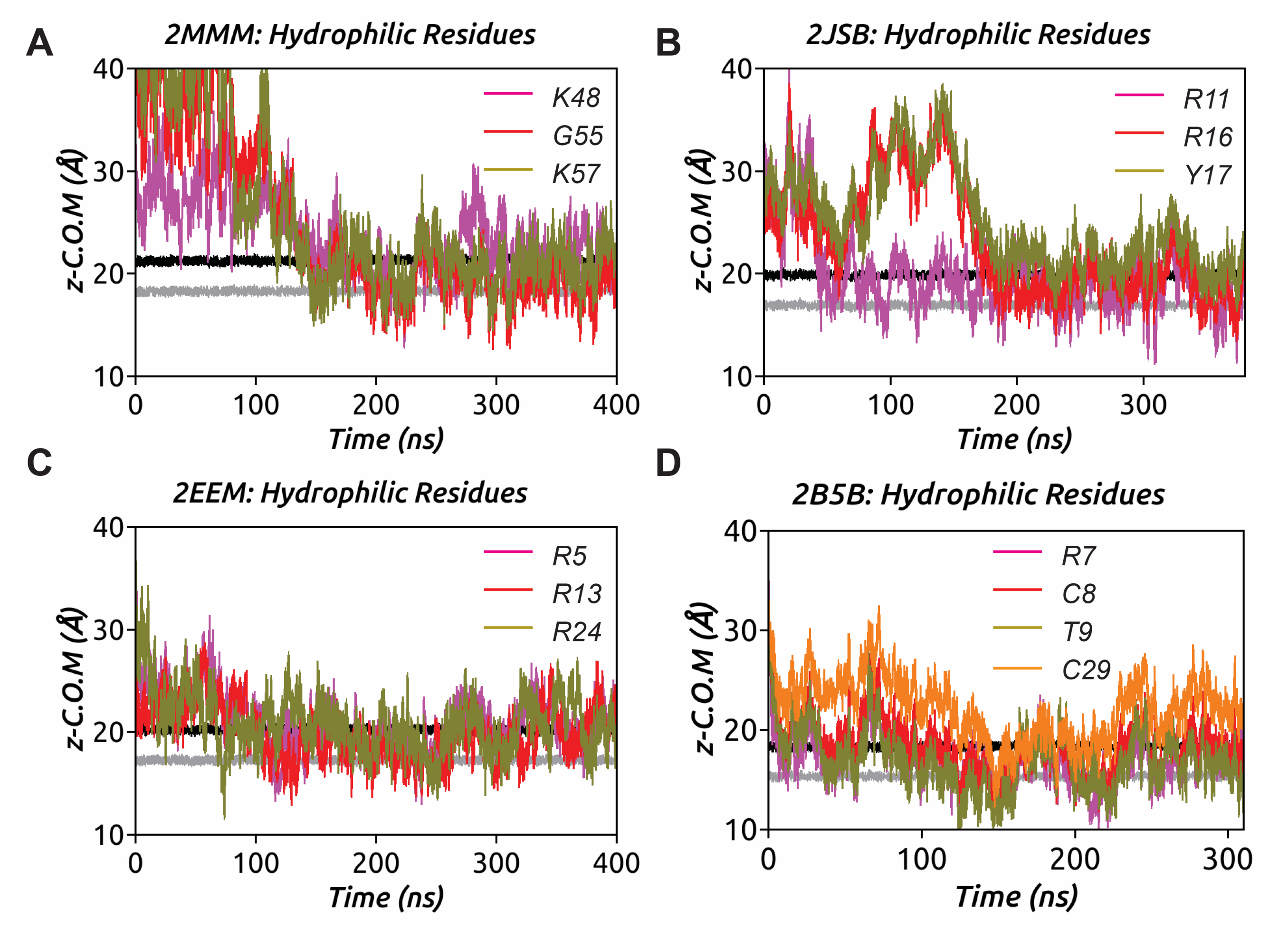


FIG. S.6. Partitioning of key hydrophilic residues into the model bacterial membrane during the simulation for four AMPs considered. The locations of headgroup and the XXX of the upper leaflet, along the membrane normal, of the model bacterial membrane are also shown in black and dark grey lines.

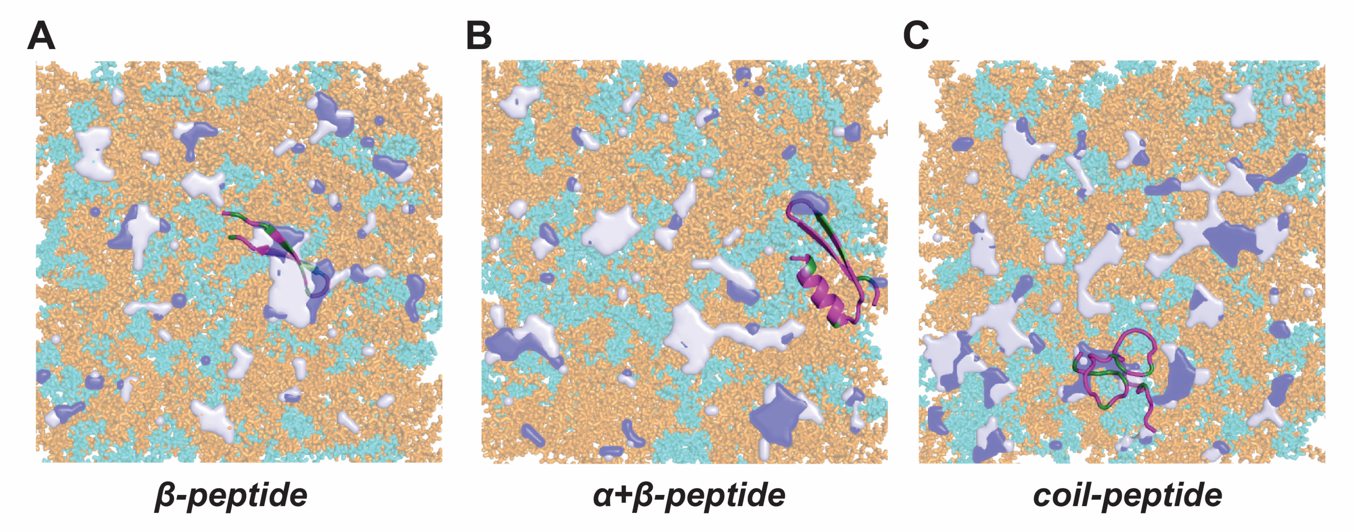


FIG. S.7. The final snapshot of AMPs-POPE (orange)/PG (cyan) system illustrating deep (dark blue) and shallow defect (light blue) sites.
